# The medial prefrontal cortex during flexible decisions: Evidence for its role in distinct working memory processes

**DOI:** 10.1101/2023.05.22.541807

**Authors:** Kevan Kidder, Ryan Gillis, Jesse Miles, Sheri Mizumori

**Affiliations:** University of Washington

**Keywords:** mPFC, spatial working memory, decision-making, behavioral flexibility, reversal learning, optogenetics

## Abstract

During decisions that involve working memory, task-related information must be encoded, maintained across delays, and retrieved. Few studies have attempted to causally disambiguate how different brain structures contribute to each of these components of working memory. In the present study, we used transient optogenetic disruptions of rat medial prefrontal cortex (mPFC) during a serial spatial reversal learning (SSRL) task to test its role in these specific working memory processes. By analyzing numerous performance metrics, we found: 1) mPFC disruption impaired performance during only the choice epoch of initial discrimination learning of the SSRL task, 2) mPFC disruption impaired performance in dissociable ways across all task epochs (delay, choice, return) during flexible decision-making, 3) mPFC disruption resulted in a reduction of the typical vicarious-trial-and-error (VTE) rate modulation that was related to changes in task demands. Taken together, these findings suggest that the mPFC plays an outsized role in working memory retrieval, becomes involved in encoding and maintenance when recent memories conflict with task demands, and enables animals to flexibly utilize working memory to update behavior as environments change.

## 1 INTRODUCTION

Memory-guided decisions are successful when prior learning matches our current expectations. When goals change, however, we must flexibly update both our behavior and the memories we use to guide our decisions. This process of updating is said to rely on working memory which involves encoding of new information, as well as maintenance and retrieval of that information to guide upcoming behavior (Baddeley, 2011; Becker et al., 1981; Buzsáki et al., 2021; Cohen et al., 1997).

Neurophysiology studies have found correlates for each of these three components of working memory (encoding, maintenance, retrieval) in prefrontal cortices. Indicative of encoding, mPFC cells respond to choice outcomes (Horst & Laubach, 2012; Luk & Wallis, 2009; Pratt & Mizumori, 2001; Y. Yang & Mailman, 2018) in tasks that require use of different strategies to earn reward. mPFC cells can also encode switches in strategy use (Hasz & Redish, 2020) and generalized task variables (Samborska et al., 2022). Together, these studies suggest that the mPFC can represent information about rewards, task structures, and rules that can be encoded for later retrieval to guide behavior.

Cells in the primate prefrontal cortex (PFC) can also exhibit elevated activity throughout the duration of a delay period, which is when information needs to be maintained in working memory to guide an upcoming decision (Batuev et al., 1979; Funahashi et al., 1989; Fuster & Alexander, 1971; Kubota & Niki, 1971). Similar delay-firing has been found in the rodent medial prefrontal cortex (mPFC) along with populations of cells whose collective activity tiles delay periods (Baeg et al., 2003; Bolkan et al., 2017; Jung et al., 1998). These results have often been interpreted as a physiological basis for active working memory maintenance (Goldman-Rakic, 1995; Zylberberg & Strowbridge, 2017).

In addition to working memory maintenance, mPFC cells have been shown to retrieve task-relevant information that guides current and upcoming choices. These cells can fire specifically at or before choice points where rats turn one direction or another in spatial delayed alternation tasks, and their firing can predict success on these tasks (Guise & Shapiro, 2017; Ito et al., 2015; Luk & Wallis, 2009; Stout & Griffin, 2020; S. T. Yang et al., 2014; Y. Yang & Mailman, 2018). Many studies also show increased hippocampal (HPC)-mPFC oscillatory coherence in the 4-12 Hz theta band surrounding choice points (Benchenane et al., 2010; Jones & Wilson, 2005; Tamura et al., 2017), suggesting choice points of tasks are times when information is retrieved to guide decision-making.

In addition to physiological studies, behavioral studies involving manipulations of the mPFC support the idea that the mPFC plays a role in working memory processing. Lesions and pharmacological inactivations decrease choice accuracy in spatial delayed alternation and non-match to place tasks, and impair switches between reward contingencies in reversal learning and set shifting tasks (Avigan et al., 2020; Birrell & Brown, 2000; De Bruin et al., 2000; Kinoshita et al., 2008; Ragozzino et al., 1999) demonstrating impairments in working memory. These studies, however, were unable to address whether distinct components of working memory (i.e. encoding, retrieval, maintenance) were affected by the mPFC manipulations because of their permanent or long-lasting effects.

Recent work using temporally precise mPFC manipulations has begun to assess causal links between disrupted mPFC function and working memory processes. Specifically, inhibiting somatostatin (SST) or parvalbumin (PV) expressing cells in the mPFC during early delay periods impaired performance on an auditory go/no-go task (Kamigaki & Dan, 2017). Another study found that either activating or suppressing mPFC activity during delays in an odor-based non-match to sample task decreased task performance when mice were learning a discrimination rule (Liu et al., 2014). Interestingly, well trained mice performed worse when mPFC activity was suppressed during the decision-making/sample matching portion of the task, but not when the mPFC was manipulated during the delay (Liu et al., 2014), suggesting the mPFC is involved in distinct working memory processes as behavior is shaped by learning.

We followed up on these reports by disrupting the mPFC at different epochs of a decision in a spatial delayed alternation (SDA) task (Kidder et al., 2021). Starting with the premise that different working memory processes were associated with different stages of the task, we reasoned that encoding began during reward delivery as rats returned to a waiting location (the return epoch), working memory maintenance occurred as rats held information about their prior choice during a delay period (delay epoch), and retrieval into working memory occurred as rats made choices about where to go for reward (choice epoch). Thus, decision-making impairments could be associated with different working memory processes based on the epoch when mPFC disruption occurred. Interestingly, we found deficits in choice accuracy when the mPFC was disrupted only during the choice epoch, but not delay or return epochs, suggesting a major role for the mPFC in working memory retrieval during spatial delayed alternation. Additionally, disrupting the mPFC during any epoch caused a decrease in a decision-making behavior known as vicarious trial and error (VTE), where decision-makers appear to vacillate between options before settling on a final decision. Furthermore, the magnitude of the decreased VTE occurrence was correlated with the magnitude of the choice accuracy deficit.

However, because the SDA task only required encoding which location was visited every trial, accurate decision-making did not depend on encoding or maintaining reward information about recent decisions. To ensure that successful performance relied on encoding and maintaining information about task-relevant variables, in this study we employed the same epoch-based disruption procedure on a serial, spatial reversal learning (SSRL) task, where reward locations switched (reversed) dynamically based on recent performance. If, as in Kidder *et al*. (2021), we assume the working memory processes are reliably separated throughout trial epochs, this design allows us to make predictions about what behavior should look like when different working memory processes are disrupted. Much like the SDA task, disrupting the mPFC during retrieval should always cause choice accuracy impairments, leading to fewer reversals and increases in the average number of trials to criterion. Because new reward contingencies need to be encoded as reversals occur, disrupting the mPFC during encoding should lead to errors caused by regression to behavior consistent with the prior contingency. Similarly, accurate decision-making should rely on maintaining information about prior choices until new contingencies are re-encoded. This implies disrupting the mPFC during working memory maintenance should also cause performance deficits on trials close to reversals, causing perseveration behavior consistent with the prior contingency.

Our results indeed show that disrupting the mPFC during each epoch can cause performance deficits. The most severe deficits were caused by disruption during the choice epoch, suggesting that the mPFC plays an outsized role in retrieving information into working memory. Furthermore, disruption during the delay epoch caused increases in the number of errors immediately following reversals without altering other performance metrics, and disruption during the return epoch caused fewer total reversals per session. What’s more, mPFC disruption, regardless of epoch, prevented the typical aggregation of vicarious trial and error behavior around reversals without decreasing their overall prevalence, suggesting a general failure to correctly time flexible behavior with respect to changes in task demands.

## 2 METHODS

### 2.1 Animals

Five Long-Evans rats (Charles River) were used in this study. The cohort consisted of 3 males (320-400 grams) and 2 females (180-220 grams). Animals were housed on a 12 hour light/dark cycle (lights on at 7:00 am) with *ad libitum* access to water. Rats were free fed upon arrival for one week, after which they were food restricted to 80-85% of their original free fed weight. Animals were only trained and tested during the light portion of their light/dark cycle. All procedures were in accordance with the University of Washington’s Institutional Animal Care and Use Committee guidelines (Protocol 3279-01).

### 2.2 Apparatus

The spatial serial reversal learning (SSRL) task took place on a fully automated plus-maze elevated 79 cm from the floor. Arms of the maze measured 58 x 5.5cm. The north and south arms were designated as start-arms, and the east and west arms were designated as goal-arms. Attached to the end of the goal-arms were 3D printed food wells connected to computer-controlled pellet delivery hardware (Med-Associates Inc.) which delivered sucrose pellets (45mg; TestDiet). The maze was remotely controlled by LabVIEW 2016 software (National Instruments) with custom built task programs. Each maze arm was hinged midway so that the proximal end could be raised and lowered via servos connected to Arduino boards. The maze was surrounded by black curtains with several visual cues attached to them so that animals could use these cues to engage spatial navigation strategies. Positioned directly above the maze was a camera (SONY) recording at ∼30Hz which integrated with LabVIEW software to identify animal location and trigger task events based on the coordinates of predetermined trigger locations.

### 2.3 Surgical Procedures and Optic Implants

Shortly after arriving at our facility, rats were anesthetized using 5% isoflurane in oxygen (flow rate 1.0 L/min) and placed into a stereotaxic apparatus (KOPF). Isoflurane concentration was then lowered to 1.0%-3.5% as rats underwent surgery involving bilateral mPFC (AP: 3.0mm, ML: ± 0.8, DV: −3.8) intracranial injections of the excitatory optogenetic viral construct AAV5-CaMKlla-hChR2-mCherry (Addgene: CS1096). 500nL of virus was injected into each hemisphere at a flow rate of 100nL/min. Following surgery, rats were allowed approximately seven days of recovery before beginning handling and maze training procedures.

Once animals reached performance criterion on the SSRL task, they underwent optic fiber implant surgery. Prior to implant surgery, optic implants were constructed using optic fiber (200µm in diameter) and ceramic ferrules held together with a quick-cure epoxy (ThorLabs). Custom implant devices which housed the two optic fibers were designed (Autodesk Inventor) and then 3D printed (Formlabs). For implant surgery, isoflurane conditions were the same as previously described, and holes were drilled into the skull at the previously used bilateral mPFC coordinates. Tips of the optic fibers were positioned just above the location of the previous viral injections (∼D/V:-3.5). Approximately 6-8 stainless steel screws were anchored into the skull, and dental repair resin (Coltene) was applied to cover the screws and hold the entire optic implant device in place. Animals were once again given approximately seven days to recover from surgery. After recovery, animals were required to meet performance criterion three consecutive days before experimental conditions began.

### 2.4 Behavioral Training and Experimental Design

#### 2.4.1 Habituation and Training

Rats were handled for 10-15 minutes on at least three occasions before they were exposed to the maze. Rats were habituated to the maze prior to behavioral training by allowing them to freely forage for sucrose pellets scattered on the maze for one session of 20 minutes. Next, animals performed a training program which consisted of 45 trials and had alternating blocks of forced choice and free choice trials in which every response was rewarded. Animals were required to complete the training program within 45 minutes for three days in a row before moving on to training on the SSRL task.

#### 2.4.2 Spatial Serial Reversal Learning (SSRL) Task

The goal of the SSRL task was for animals to disambiguate which of the two goal locations (east or west) was the current block’s correct (reward) location. At the start of each session the initial correct arm was randomly set by the experimenter and each block ended when the animal chose the correct arm 9/10 times. At the beginning of each new block the reward location was switched to the opposite reward arm (Figure 1a). Animals were then tested to see how many reward arm reversals they could complete within a session. The first block of a session was considered the initial discrimination (ID) block, and all subsequent blocks were considered reversal blocks.

**Figure 1.**
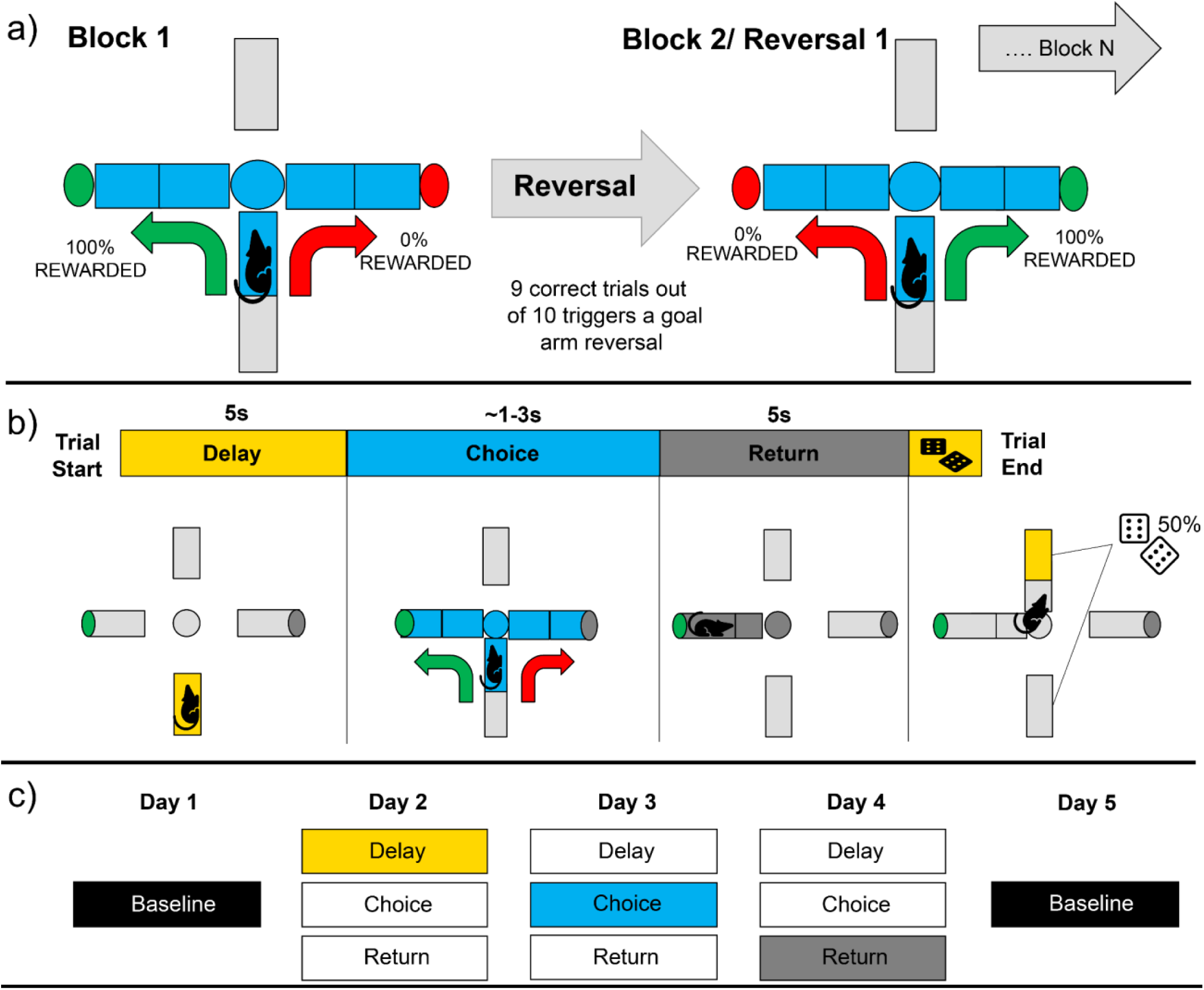
Illustration of the spatial serial reversal learning (SSRL) task. a) Animals underwent five days of experimental sessions. Animals started with a baseline session when no optogenetic disruption occurred, and in the subsequent three days the animal experienced optogenetic mPFC disruption in a selected epoch for every trial in the session (stimulated epoch order was randomized across animals). Experimental sessions concluded with a final baseline day when no mPFC disruption occurred. b) Animals first showed discrimination of the initially rewarded arm by selecting the correct arm 9 out of 10 times. After this the reward location was reversed to the opposite reward arm. c) depiction of SSRL task epochs (delay, choice, return).

A single session of the SSRL task consisted of 200 trials and each trial consisted of three epochs: delay, choice, and return (Figure 1b). At the start of each trial, all maze arms were lowered so that the animal was restricted to their current start-arm where they waited through the five second delay epoch. After the delay, the current trial’s start-arm and both goal-arms were raised so that the animal could navigate to the goal location of their choice. A typical choice epoch, which started by raising the start arm after the delay and ended when an animal reached a goal location, lasted 1.5-3 seconds. After reaching a goal location, animals received one sucrose pellet for correct responses and no sucrose pellets for incorrect responses. During this time, the next trial’s start-arm was randomly chosen and subsequently raised so that the animal could travel there and trigger the start of the next trial’s delay. Therefore, the return epoch was defined from the point of the animal reaching a goal location, included consuming (or not consuming reward), and ended when the animal reached the next trial’s randomly chosen start-arm. Return epochs generally lasted five seconds. Animals reached performance criterion on the SSRL task once they reached asymptotic performance which was measured as the total number of reversals completed in a session. After reaching the initial SSRL performance criterion animals underwent optic fiber implantation surgery.

#### 2.4.3 Experimental Design

Each animal underwent a total of five experimental conditions which occurred in separate sessions, each on a separate day (Figure 1c). These conditions included: (1) a pre-stimulation assessment, (2-4) mPFC optogenetic disruption during each of the three task epochs (delay, choice, return; order of epoch-specific disruption sessions was counterbalanced for each animal), and (5) a post-stimulation assessment. There were no significant differences between pre- and post-stimulation assessments, indicating no lasting effects of optogenetic stimulation (Figure S1). Therefore, the average of pre- and post-stimulation assessment sessions were combined to form the baseline for comparison in subsequent data analyses.

#### 2.4.4 Optogenetic Stimulation

To activate opsins and disrupt mPFC neural activity, a 473nm laser (Laserglow Technologies) was held to a power of ∼6-7mw emanating from the tip of the implanted optic fibers (measured prior to optic implant surgery). Stimulation was delivered at a rate of 20 Hz with a 50% duty cycle for the duration of the selected epoch (delay, choice, or return). Stimulation rate was controlled by an Arduino which also received signals from the LabVIEW program to gate laser activation according to the specific epoch being tested. A single fiber optic cable was connected from the power source to an optic commutator (Doric) mounted above the maze, from which two fiber optic cables were affixed. These fiber optic cables were then connected to the animal’s optic ferrules with copper sleeve connectors (ThorLabs).

### 2.5 Histology

After all conditions were completed rats were given an overdose of sodium pentobarbital. Once rats were deeply anesthetized, they were transcardially perfused with 0.9% saline and 10% formaldehyde solution. Brains were extracted and stored at 4°C in formalin for a day and then submerged in a 30% sucrose solution for four days. Brains were then frozen and cut into coronal sections (45 μm) on a freezing microtome. Brain slices were then mounted onto slides and fluorescence was preserved with the mounting medium Vectashield (Vector Laboratories). Slices were examined with a fluorescent microscope to verify viral expression and optic fiber placement in the mPFC. Only animals with proper bilateral viral expression and optic fiber placement contributed to subsequent data analysis.

### 2.6 VTE Identification and Analysis

Using the same videos recorded during in-task position tracking (see **Apparatus** above), we applied DeepLabCut (DLC) version 2.2 to identify the heads of implanted rats running the SSRL task. We initially labeled 20 or 30 frames for each session using the built-in labeling GUI for a total of approximately 400 labeled frames for the first model training attempt. Because many of our images were low contrast and mislabeled, we relabeled 20-50 outliers in a subset of videos and retrained a new iteration of the network on the now expanded dataset to achieve satisfactory performance. Each training attempt used NVIDIA GEFORCE GTX 1080 GPU with 500,000 iterations.

Choice epoch segmentation for trajectory detection was accomplished by detecting when rats crossed the maze central platform, then moving earlier in the trajectory to a user defined starting point and later in the trajectory to a user defined ending point. Before epoch segmentation, all position data from DLC were rotated and scaled into maze coordinates with the center of the maze at approximately the origin of an (x, y) grid. Position data were median filtered with a 7-point window to mitigate any jumps from DLC tracking and all choice epoch trajectories were quality checked by experimenters.

Trajectories with vicarious trial and error (VTE) were detected by projecting the position data into principal component (PC) space and clustering the PC- representations of the trajectories with hierarchical agglomerative clustering. Before projection, all trajectories were aligned and standardized to the same starting and ending positions. Visual inspection of clustering in PC space naturally formed what looked like two clouds in low dimensional plots, and distance-based dendrograms cut to give two clusters separated trajectories with VTE from non-VTE trajectories.

## 3 RESULTS

### 3.1 Histology

Tips of optic fibers were located bilaterally in the mPFC (Figure 2a). Optic fiber tip placements were evenly distributed along the D/V axis of the prelimbic cortex, ranging from just below the ACC-PL border to the PL-IL border. Viral expression in the mPFC surrounded all optic fiber tips, and expression was relatively equal between mPFC hemispheres. Animals without proper optic fiber placement or viral expression were not included in the data analysis.

**Figure 2.**
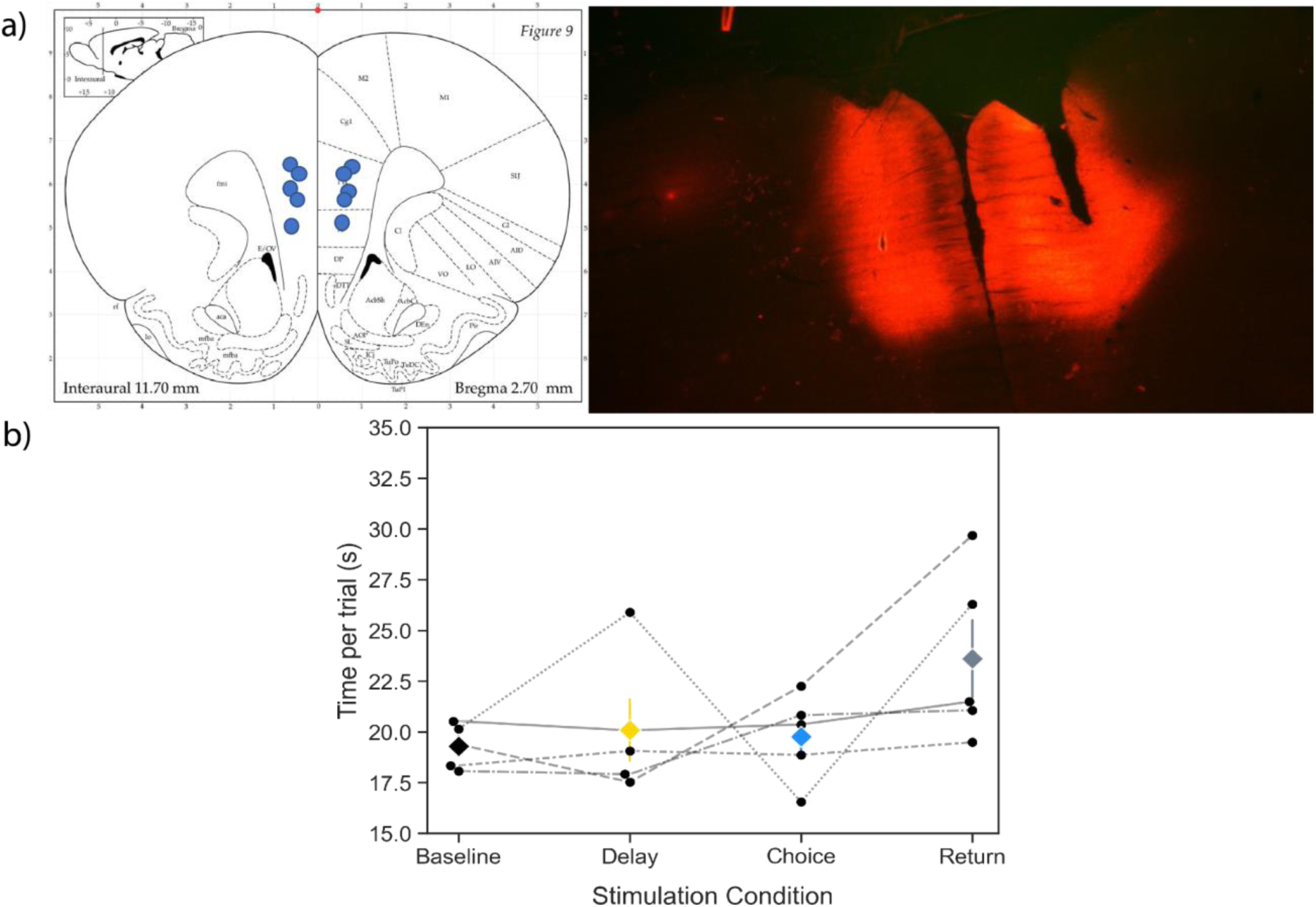
a) mPFC optic fiber tips terminated within the prelimbic/infralimbic division of the mPFC. Only animals with clear bilateral mPFC expression were used for analysis. b) Time per trial was not significantly different (*p* > .05) between conditions, suggesting stimulation did not impair animals’ motivation to complete the task.

### 3.2 mPFC disruption impairs SSRL performance

To validate that the mPFC is involved in this task, we combined all stimulation conditions together and compared the distributions of average trials per block for stimulation (x̄ = 18.8, SE = 1.05) and baseline conditions (x̄ = 13.9, SE = 0.47) (Figure 3a). A t-test revealed the distributions are significantly different from each other (t = − 6.89, *p*<0.05). These data reveal that regardless of which epoch disruption was applied to, mPFC disruption impaired performance on the SSRL task. Additionally, time per trial was not significantly different between baseline or stimulation conditions (t = −1.12, p =0.277) suggesting mPFC disruption did not alter animals’ motivation to engage in the task (Figure 2b).

**Figure 3.**
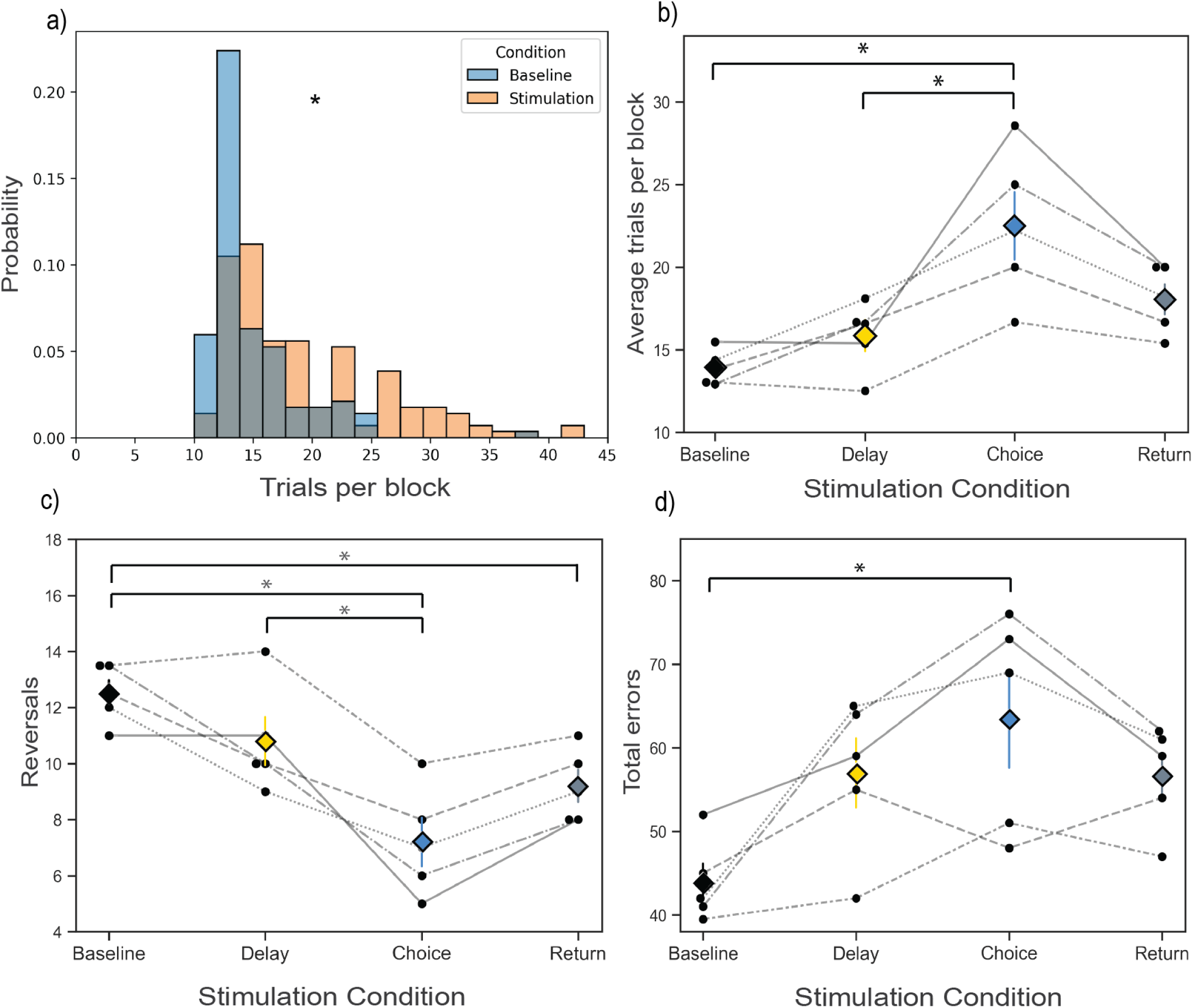
a) There was a significant difference (\**p*<0.5) in the probability of trials per block between baseline and stimulated sessions. b) Choice epoch disruption significantly increased the trials in a block compared to delay epoch disruption and baseline. c) Choice epoch disruption significantly reduced the number of reversals completed in a session compared to delay epoch disruption, and choice and return epoch disruption significantly reduced the number of reversals completed in a session compared to baseline. d) Choice epoch disruption significantly increased the total number of errors compared to baseline.

### 3.3 Differential epoch-specific effects on SSRL performance

To determine if mPFC disruption during a particular epoch (delay, choice, return) selectively impaired performance, we compared the average trials per block, average number of reversals, and total errors across baseline and the three epoch stimulation conditions (Figure 3b-d). There was an effect of stimulation epoch on the average trials per block (F(3) = 15.67, p = 0.0002) in which choice epoch stimulation caused significantly more trials per block (x̄ = 22.49, SE = 2.04) than baseline (x̄ = 13.92, SE = 0.47) and delay stimulation (x̄ = 15.85, SE = 0.94), but not return stimulation (x̄ = 18.04, SE = 0.91) (Figure 3b). A repeated measures ANOVA revealed an effect of stimulation epoch on the average number of reversals (F(3) =23.93, p<0.0001). Post-hoc tests for multiple comparisons revealed the average number of reversals in the choice epoch stimulation condition (x̄ = 7.20, SE = 0.86) was significantly less than baseline (x̄ = 12.5, SE = 0.47) and delay epoch stimulation (x̄ = 10.80, SE = 0.86), but was not different from return stimulation (x̄ = 9.20, SE = 0.58), which also showed significantly less reversals than baseline (Figure 3c). There was a significant stimulation epoch effect on the total number of errors (F(3) = 9.02, p = 0.0021): stimulating during the choice epoch caused significantly more errors (x̄ = 63.4, SE = 5.8) than baseline (x̄ = 43.9, SE = 2.21), but not delay (x̄ = 57.0, SE = 4.16) or return stimulation (x̄ = 56.6, SE = 2.77) (Figure 3d). To summarize, stimulation across epochs tended to impair performance. However, choice epoch stimulation was the only epoch in which performance was consistently and significantly impaired. Return epoch stimulation significantly impaired performance only by reducing the total numbers of reversals in a session, while delay epoch stimulation did not significantly impair performance on any of these metrics.

### 3.4 mPFC choice epoch disruption impairs both initial discrimination and reversal performance

While there is consensus regarding the mPFC’s involvement in discriminating between similar recent memory representations (goal locations) during reversal blocks of similar tasks, prior work (Avigan et al., 2020; Guise & Shapiro, 2017) has shown conflicting results regarding the mPFCs role in initial discrimination learning. We therefore analyzed the effects of mPFC disruption between ID and reversal blocks of this task (Figure 4). There was an effect of stimulation epoch on both the number of trials to ID (F(3) = 3.97, p= 0.035) and trials per reversal (F(3)= 17.42, p=0.0001). In both cases, choice epoch stimulation increased the average number of trials (ID: x̄ = 25.0, SE = 2.98; reversal: x̄ = 27.41, SE = 1.23) compared to baseline (ID: x̄ = 15.80, SE = 0.93; reversal: x̄ = 15.50, SE = 0.26) (Figure 4a-b). We next analyzed the occurrence of errors between ID and reversal blocks. There was an effect of stimulation epoch on both the number of ID errors (F(3) = 3.35, p=0.056) and average reversal errors (F(3) = 11.39, p= 0.0008) (Figure 4c-d). In both cases, choice epoch stimulation increased the number of errors committed (ID: x̄ = 6.6, SE = 1.21, reversal: x̄ = 7.62, SE = 1.23) compared to baseline (ID: x̄ = 2.8, SE = 0.34, reversal: x̄ = 3.12, SE = 0.26). Lastly, repeated measures ANOVAs revealed no effect of stimulation condition on ID accuracy (F(3) = 1.52, p=0.26) but did reveal an effect on reversal accuracy (F(3) = 9.40, p= 0.0018) (Figure 4e-f). Choice epoch stimulation decreased reversal accuracy (x̄ = 0.59, SE = 0.033) compared to baseline reversal accuracy (x̄ = 0.72, SE = 0.014) but was not different from return (x̄ = 0.64, SE = 0.016) or delay (x̄ = 0.65, SE = 0.021) which were both not significantly different from baseline.

**Figure 4.**
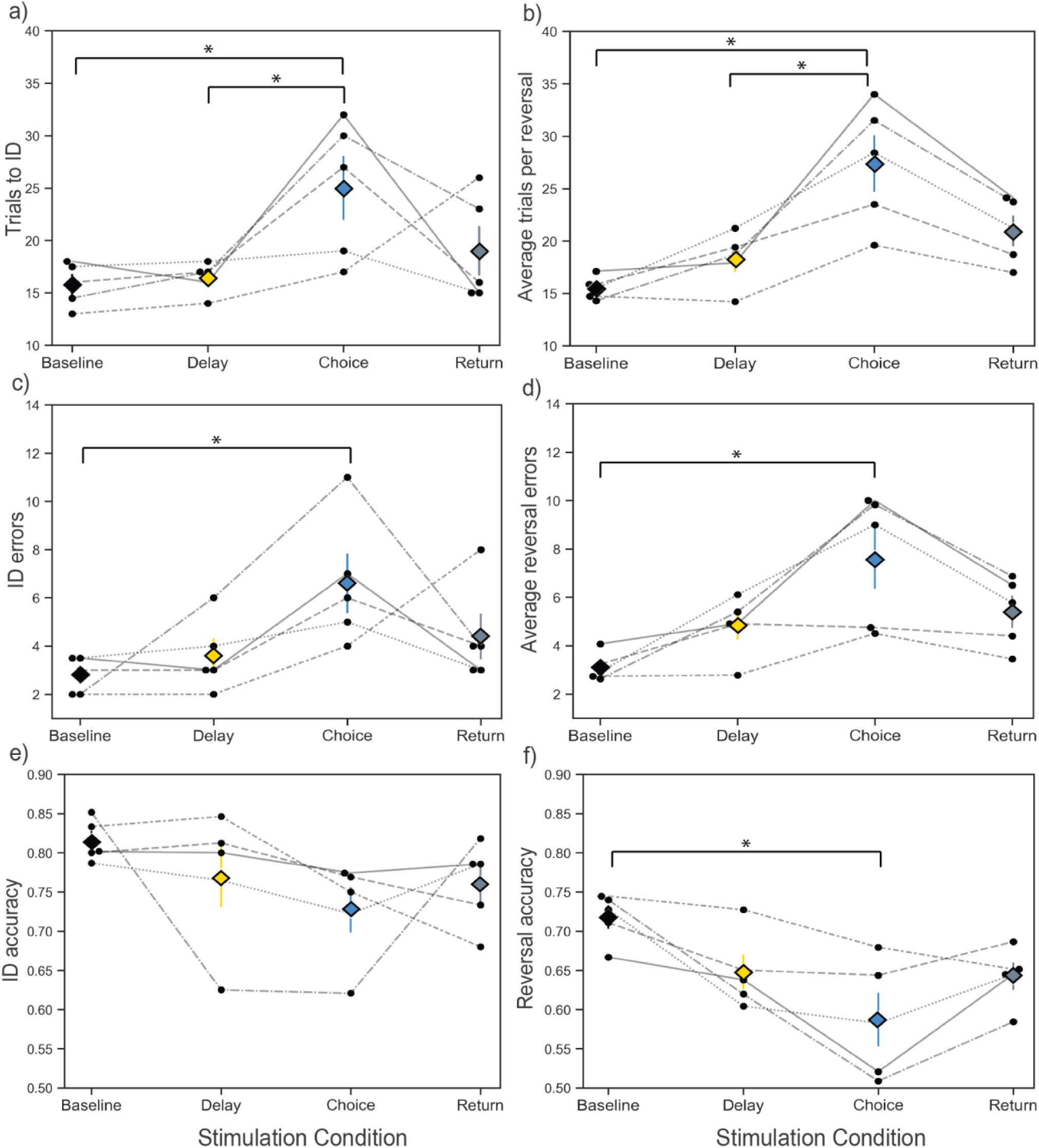
Choice epoch disruption significantly increased (\**p*<.05) both the number of trials to initial discrimination (ID) (a) and the average trials per reversal (b). Choice epoch disruption significantly impaired the number of errors in ID (c) and reversal (d) blocks. ID choice accuracy (e) was not impaired by mPFC disruption in any condition, however choice disruption did significantly impair reversal blocks choice accuracy (f). Therefore, the only difference in effect of mPFC stimulation occurred in terms of choice accuracy between ID and reversal blocks.

These results reveal mPFC stimulation impaired both ID and reversal performance to some extent. However, while impacts to ID and reversal performance were similar across most metrics, reversal performance was impacted to a greater extent as revealed by a selective decrease in choice accuracy in reversal blocks and not ID blocks following choice epoch disruption.

### 3.5 Perseverative and regressive errors reveal dissociable working memory roles for the mPFC

We identified perseverative errors as all errors that occurred before the animal made two correct choices in a new block, while regressive errors were those that occurred after the animal made two correct choices in the new block. There was an effect of stimulation epoch on both the number of perseverative (F(3) = 7.50, p= 0.0044) and regressive (F(3) = 9.63, p= 0.0016) errors (Figure 5a-b). Animals committed significantly more perseverative errors during the delay epoch stimulation condition (x̄ = 24.4, SE = 2.36), compared to baseline (x̄ = 15.6, SE = 1.76), while errors following choice (x̄ = 16.8, SE = 1.24) and return (x̄ = 16.8, SE = 1.83) stimulation were not significantly different from baseline. Conversely, there were significantly more regressive errors in the choice epoch condition (x̄ = 33.6, SE = 5.35) than baseline (x̄ = 14.2, SE = 2.19) and delay (x̄ = 18.2, SE = 3.04) conditions, but not return condition (x̄ = 26.2, SE = 2.22).

**Figure 5.**
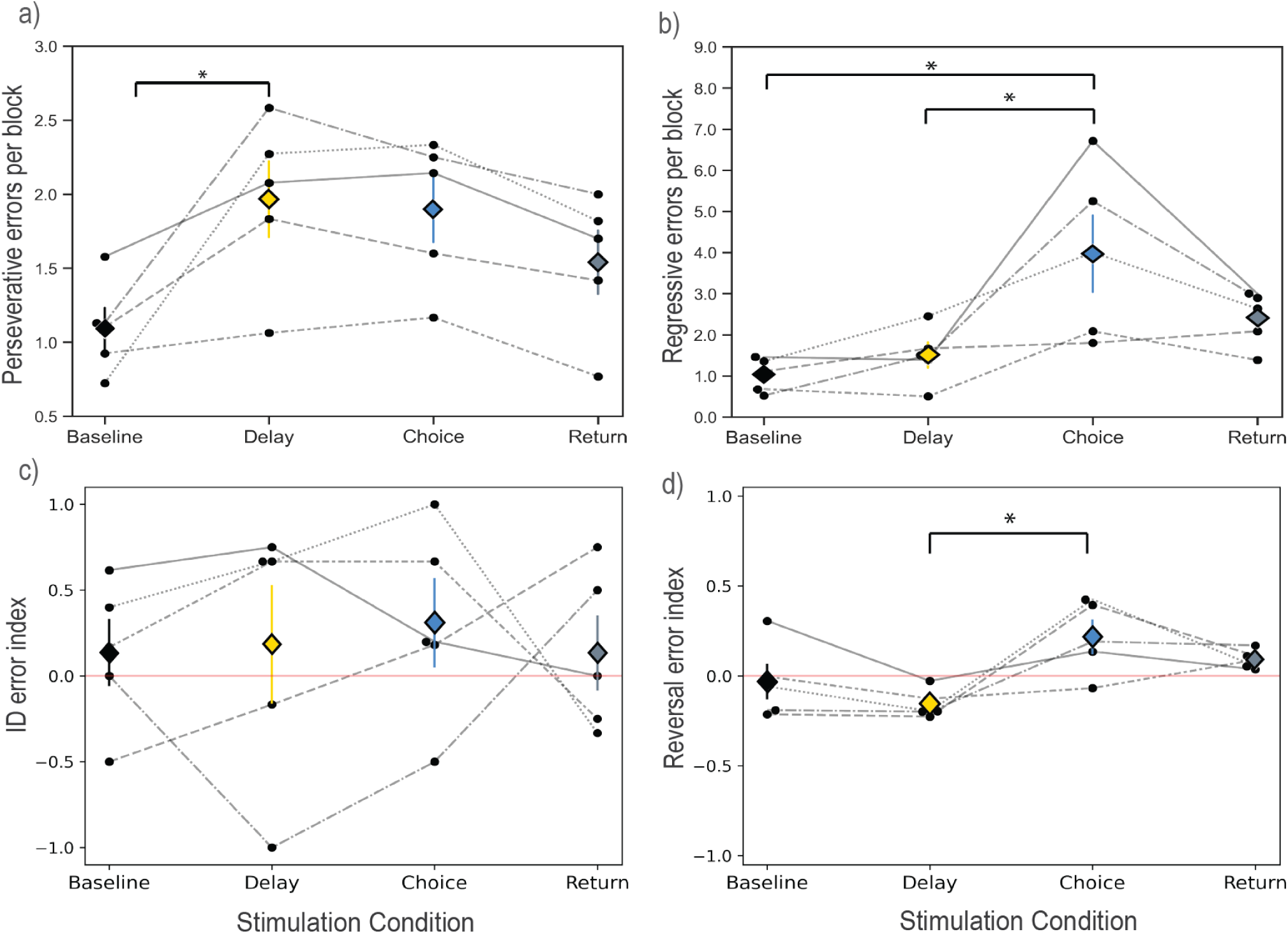
After a reversal, perseverative errors were defined as all errors that occur before the animal makes 2 correct choices in a row. Perseverative errors are therefore thought to reflect an inability to learn the new strategy. After the animal makes two correct choices in a row all subsequent errors in the block are defined as regressive errors. Regressive errors are thought to reflect a failure to maintain or retrieve the new strategy. A positive error index means that more regressive errors than perseverative errors occurred in a block, while a negative error index means that more perseverative errors occurred than regressive errors in a block. a) Delay epoch disruption significantly increased (\**p*<.05) perseverative errors compared to baseline. b) Choice epoch disruption significantly increased regressive errors above baseline and delay epoch disruption. c) There were no significant differences between average error index values by stimulation condition in the ID block. d) Average error index values in reversal blocks for the delay and choice disruption groups were significantly different.

We then created an error index metric which involves a ratio of perseverative to regressive errors in order to see how the distribution of regressive and perseverative errors in individual blocks changed across task and stimulation conditions. If more regressive errors than perseverative errors occurred in a block, the error index value is positive. The error index value for a block is negative if more perseverative errors than regressive errors occurred in that block. During the ID there were no significant differences between the error index values of stimulation epoch conditions (F(3) = 0.11, p=0.96) (Figure 5c). Furthermore, all stimulation conditions contained slightly positive error index values, indicating there were always more regressive errors during ID blocks. During reversal blocks (F(3) = 4.71, p=0.02) however, delay epoch stimulation resulted in a switch from a positive to a negative error index value, and this was significantly different from the choice epoch stimulation condition (p=.007) (Figure 5d). These data reveal that mPFC disruption during reversal blocks, not ID blocks, biased the types of errors that occur in a block depending on which epoch the mPFC disruption occurred.

### 3.6 VTE rate fluctuates according to task demands and is impaired by mPFC disruption

Out of the 3561 trials analyzed, 686 (19.3%) were identified as VTE trials. Unlike Kidder et al. (2021), we do not see differences in VTE rates per block when we disrupt the mPFC (t = 0.621, p = 0.535, Figure 6b). Instead, mPFC stimulation altered the dynamics of VTE occurrence surrounding reversals. During baseline sessions, the average rate of VTE occurrence is higher than average early in the block, while near the end of a block the rate of VTE occurrence is lower than average (Figure 6b, top-center). Furthermore, the rates appear periodic, showing regularly spaced peaks in the average reversal-aligned VTE-rate autocorrelation functions (Figure 6b, top-right). In contrast, sessions with stimulation do not tend to show the same regular increases and decreases in VTE rate aligned to reversals. Rather, these sessions tend to show the same average VTE rate (approximately 20% of trials) regardless of proximity to a reversal. This is further indicated by the almost entirely flat autocorrelation functions for reversal-aligned VTE rates from stimulation sessions.

**Figure 6.**
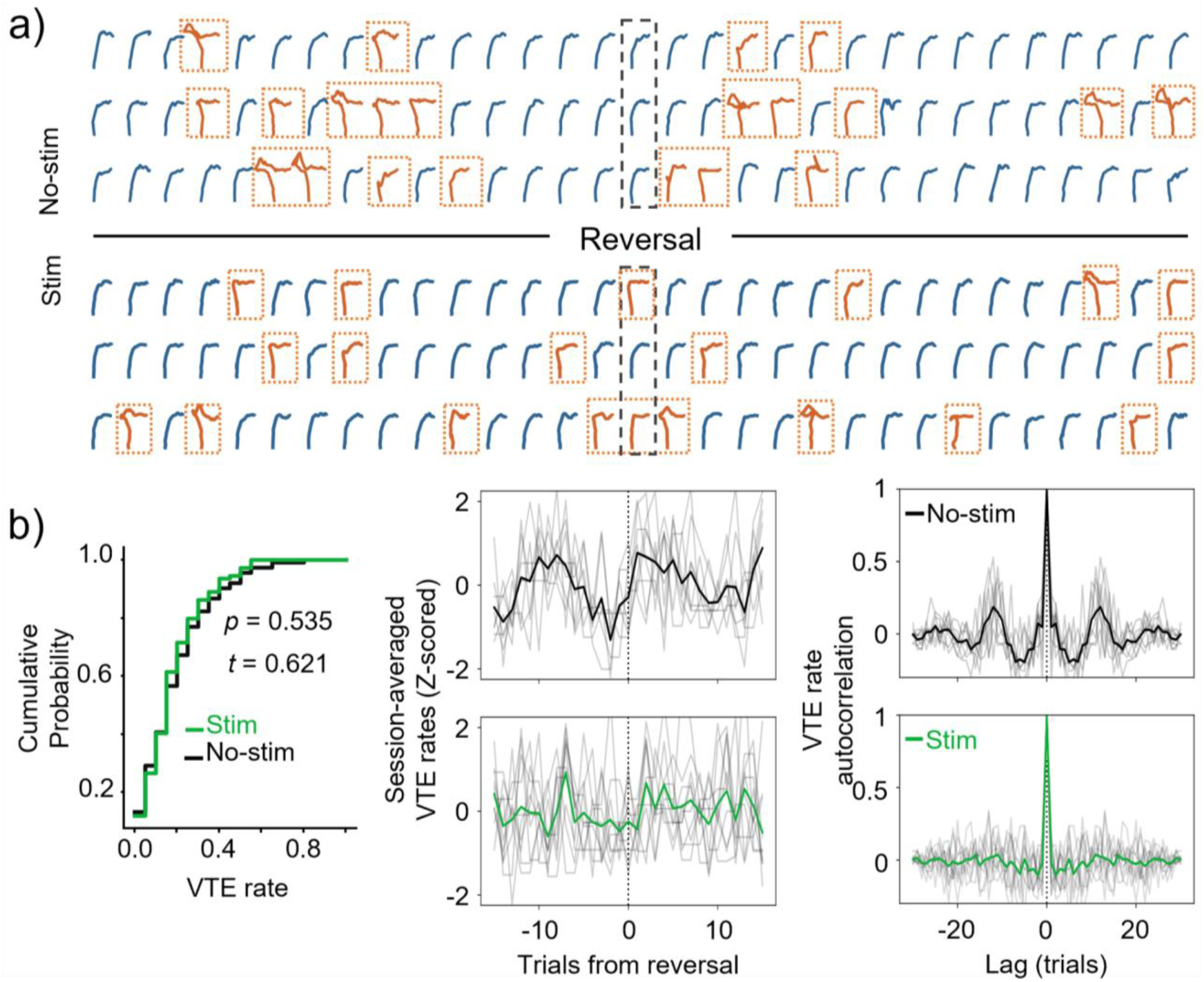
Coordination of vicarious trial and error (VTE) behavior with task demand is sensitive to mPFC disruption. The top of panel a) shows sequences of choice trajectories from 15, non-stimulated trials before (left of dashed box) and after reversals (right of dashed box) for three different reversal blocks. Choice trajectories showing VTE are colored orange and surrounded by a dotted box. Choice trajectories showing VTE for stimulation trails are shown below the black line. All trajectories are oriented so they start at the bottom and end at the right, and thus do not reflect their actual starting and ending points. In b), the left panel shows cumulative probability density functions for VTE rates (calculated on reversal blocks) during stimulated sessions in green and non-stimulated sessions in black. There is no significant difference in the distribution of block-averaged VTE rates between stim and no-stim conditions (*p* = 0.535, *t* = 0.621, two-tailed, two-sample T-test). The middle panel shows session-averaged VTE rates aligned to reversals, with non-stimulated sessions on top (black) and stimulated sessions on bottom (green). Thin, gray lines are individual session averages (across reversal blocks) and bold, colored lines are the averages across individual sessions. On the right are autocorrelation functions for each respective VTE rate time-series from the middle panel. Note the clear periodicity in VTE rates as a function of trials from reversal when no stim is applied, and general adherence to mean rates when the mPFC is stimulated.

## 4 DISCUSSION

Findings from this study provide direct causal evidence of mPFC involvement, albeit in different ways, in distinct working memory processes during flexible decision-making in the SSRL task. Consistent with past studies, we found that bilateral mPFC optogenetic disruption impaired overall performance on the SSRL task. Furthermore, selectively, and separately disrupting the mPFC in each task epoch (delay, choice, return) impaired performance during reversal blocks, while mPFC disruption during initial discrimination (ID) blocks impaired performance only when the disruption occurred within the choice epoch. An analysis of perseverative and regressive errors during reversal blocks revealed the mPFC is critically and differentially involved during the delay and choice epochs of our task. Specifically, this error analysis revealed that when the mPFC was disrupted during delay epochs, animals tended to maintain the previous strategy longer (i.e. showed more perseverative errors), while mPFC disruption during choice epochs tended to cause a failure to retrieve or implement the newly learned strategy (i.e. showed more regressive errors). Lastly, our analysis of VTE behavior showed that disrupting the mPFC during the SSRL task does not impair the average rate of VTE behavior. Instead, it altered the dynamic fluctuation in VTE rates around changes in task demands. Together, these findings reveal differential mPFC involvement in the three working memory processes, and a role for the mPFC in coordinating deliberative behaviors during flexible memory-guided decision-making.

### 4.1 mPFC is involved in working memory encoding, maintenance, and retrieval during flexible decision-making

Past research on the mPFC suggests it has an active role in the encoding, maintenance, and retrieval of task relevant features in working memory (Luk & Wallis, 2009; S. T. Yang et al., 2014). Evidence for the mPFC’s role in working memory encoding comes from studies that report mPFC neurons which selectively fire in response to reward outcomes and stimulus presentations (Horst & Laubach, 2012; Warden & Miller, 2010; S. T. Yang et al., 2014; Y. Yang & Mailman, 2018). In support of the notion that the mPFC may shift roles amongst distinct working memory processes, Lundqvist et al. (2016) found increases in the rate of mPFC gamma bursts during stimulus presentations while finding both a decrease in gamma bursts and increase in beta bursts during delay periods. This shift from gamma bursts to beta bursts may be reflective of the mPFC switching between the processes of working memory encoding during reward periods to working memory maintenance during delay periods (Jayachandran et al., 2022; Lundqvist et al., 2016). Also during delay periods, neuronal responses that are selective to spatial locations and choice outcomes can be seen with persistent increased activity (Bolkan et al., 2017; Liu et al., 2014), which further suggests the mPFC maintains information in working memory across delays periods. A role for the mPFC in working memory retrieval comes from the idea that choice points of tasks are times when information must be retrieved and shared across structures to guide action selection (i.e., choice): studies report increased firing rates in mPFC single-units surrounding choice points and LFP recordings show mPFC theta power peaks during choice points (Luk & Wallis, 2009; S. T. Yang et al., 2014; Y. Yang & Mailman, 2018). Additionally, HPC-mPFC oscillatory coherence in the theta band peaks during choice points (Benchenane et al., 2010; Griffin, 2021; Jones & Wilson, 2005; Tamura et al., 2017), suggesting that choice points are critical times when the mPFC communicates with the HPC, possibly to retrieve and share memories which guide action-selection. In further support of this idea, studies (Guise & Shapiro, 2017; Schmidt et al., 2019) report mPFC pharmacological inactivation impairs spatial working memory performance through a disruption of pattern separation by hippocampal prospective codes. This result indicates that the mPFC is necessary to help inform hippocampal representations about task-relevant information which guides prospective decisions. Altogether, these studies indicate that during decision-making, the mPFC encodes task relevant features into working memory, and maintains them over delay periods to help the hippocampus engage in context appropriate memory retrieval during deliberation.

In the current study, we reasoned that the three working memory processes (encoding, maintenance, retrieval) would be differentially distributed throughout epochs of a trial. The delay epoch of the SSRL task is the time when task-relevant features need to be maintained in working memory so they can be used to guide decisions in future trials. Disrupting the mPFC during the delay epoch resulted in the least overall performance impairment and did not impair performance during discrimination learning. The lack of an effect from delay epoch disruption during ID blocks suggests the mPFC is not required for maintaining reward location in working memory when there are no conflicting recent memories (i.e. before reversals). We then conducted an error analysis during reversal blocks and revealed a slight increase in the overall number of perseverative errors that was only significant when the mPFC was disrupted during the delay epoch. This selective increase in perseverative errors implies it was harder for animals to maintain new information about reward contingencies in working memory during delay epoch disruption, subsequently leading to difficulty learning the new strategy. Interestingly, delay epoch disruption was the only condition which switched the sign of the error index value between ID and reversal blocks, indicating a stimulation-driven redistribution from regressive to perseverative errors after the ID. This pattern was significantly different from the choice epoch error index value. These findings provide direct behavioral evidence that the mPFC is involved in an active maintenance of task-representations (working memory maintenance) only once flexible decision-making is required.

The choice epoch of our SSRL task may represent the time when task-relevant information must be retrieved across the hippocampal-mPFC memory system to aid deliberation and choice selection. Selectively disrupting the mPFC during choice epochs consistently impaired performance across more metrics than any other epoch. Also, choice epoch disruption was the only condition which resulted in impairments both to discrimination learning (ID blocks) and to flexible decision-making (reversal blocks). Lastly, our analysis of errors during reversal blocks revealed a striking increase in regressive errors but not perseverative errors during mPFC choice epoch disruption. This selective increase in regressive errors implies animals had a harder time implementing the new strategy, suggesting a failure to retrieve memories related to the new strategy during mPFC choice epoch disruption. Conversely, the lack of an effect on perseverative errors while disrupting during the choice epoch suggests animals had no problem encoding the new strategy.

The return epoch of the SSRL task began at reward delivery and included reward consumption. Therefore, it included the time when outcome information was first available for encoding. Since return epoch disruption did not impair ID performance, the hippocampus was likely able to encode the initial goal location without the mPFC. This is consistent with studies showing cells in the intermediate hippocampus encode goal locations (Aoki et al., 2019; Pfeiffer, 2022) and place cells in dorsal hippocampus increase firing rates during paths towards goals (Jarzebowski et al., 2022; Tryon et al., 2017). However, our data shows that return epoch disruption did significantly impaired the total number of reversals per session. The lack of an effect from disruption during ID blocks when compared to the decreased number of reversals suggests that animals were impaired in their ability to encode new reward information during reversal blocks. This may have reduced the ability to override previously used strategies during return epoch disruption.

Interestingly, for almost all other behavioral metrics, impairments caused by return epoch disruption tended to show an effect intermediate of those seen from delay and choice epoch disruption. This may be in part because, on the SSRL task, impairments caused by a failure to encode information into working memory could, in theory, manifest in the same way as a failure to retrieve or maintain information in working memory. This is because there would be no memory to maintain or retrieve if it was never encoded in the first place. Another possibility is that our epoch designations may contain some overlap between working memory processes. For example, the return epoch of our task includes a traversal from the reward location to the start platform, which could be a time when information needs to be maintained in working memory. However, if the latter was the case we would expect to see performance deficits resulting from return epoch disruption to be more similar to those caused by delay epoch disruption. Our data however show the opposite: return epoch disruption effects are closer in magnitude to those caused by choice epoch disruption. Thus, it does not appear that there is meaningful overlap of working memory processes due to our epoch designations.

The delay epoch of the SSRL task is the time when information must be maintained in working memory. Much like our previous study (Kidder et al., 2021), we did not see any effects of delay epoch mPFC disruption during ID blocks. We reasoned that the mPFC may not have been involved during the delay epoch of the SDA task because it did not require animals to maintain outcome information in working memory. Since the present study’s SSRL task did require animals to know which location was rewarded, the lack of an effect from mPFC disruption during delay epochs, specifically in ID blocks, reveals the mPFC is not necessary to maintain spatial or outcome information in working memory when there are no recent conflicting memories.

Our results suggest that the mPFC performs working memory retrieval and/or action selection during basic discrimination learning, which is a time when only one memory representation (goal location) of the task is present. During reversal blocks, a time when multiple recent memories must be considered, mPFC disruption in each epoch impaired performance in different ways. The selective increase in regressive errors during choice epoch disruption suggests the mPFC is important for retrieving recent memories to make decisions. The selective increase in perseverative errors resulting from delay epoch disruption suggests the mPFC has a role in maintaining current strategies during delays only when task representations require updating. Lastly, impairment to the number of reversals completed as a result of return epoch disruption suggests the mPFC supports the encoding of reward information when recent memory conflicts with current task demands. Therefore, these findings reveal the mPFC becomes critically involved in all working memory processes when animals face decisions requiring the flexible use of memory to discriminate between or update conflicting memories.

### 4.2 The mPFC engages deliberative behavior according to task demands

Vicarious-trial-and-error (VTE) is a behavior in which animals vacillate between path options, presumably in an act of deliberation (Redish, 2016; Schmidt et al., 2013). In Kidder et al. (2021) we found that mPFC disruption, regardless of epoch, decreased the occurrence of VTEs. Furthermore, the decrease in the occurrence of VTEs significantly correlated with the decrease in performance (choice accuracy) caused by mPFC disruption. This demonstrated that 1) the mPFC modulates VTE behavior, and 2) VTEs are correlated with choice behavior. Numerous other studies also demonstrate VTE behavior is dependent on the mPFC and its functional interaction with dorsal hippocampus (McLaughlin et al., 2021; Papale et al., 2016; Schmidt et al., 2019).

One model suggests VTEs may act as a compensatory mechanism to aid deliberation in the face of uncertainty (Amemiya & Redish, 2016; Papale et al., 2012). In our task, uncertainty increases when recent memory conflicts with current task demands (i.e. at reversals), which is when we saw transient increases in VTE rates. Interestingly, in contrast with our previous findings (Kidder et al., 2021), the overall prevalence of VTEs did not change between baseline and stimulation conditions. This could be due to the fact that, during the SDA task, success is only determined by which reward location was immediately visited, not whether it was rewarded, and where the reward contingency never changes, resulting in constant certainty levels. In other words, by introducing periods of uncertainty into the task, we see that VTE rates do not go down when we disrupt the mPFC. Instead, disrupting the mPFC during the SSRL task decouples VTE rates from periods of task-related uncertainty. Thus, in addition to providing further support that the mPFC is causally involved in regulating VTE rates, our current results suggest that a critical role of the mPFC is adjusting when deliberative behaviors occur with respect to changes in uncertainty. Importantly, this explanation does not require undisturbed mPFC involvement for setting the overall VTE rate, which could be a function of uncertainty calculations done elsewhere in the brain (Fiorillo et al., 2003).

Recent results from Mclaughlin and Redish (2023) are also consistent with the interpretation that the mPFC is important for regulating VTE rates with respect to changing uncertainty. The authors show that optogenetically disrupting the mPFC during choices in a spatial delayed discounting task decreases VTE rates, and that these decreases were specific to trials that were close to when rats chose to stop adjusting delay durations for a larger reward. Control rats generally showed higher VTE rates during this period than stimulated rats. Taken together with their other results, the authors use these decreased VTE rates during mPFC disruption to suggest that the mPFC is important for flexibly updating decision-making from deliberative to procedural strategies, which could be another way of describing the decrease in VTE rates that we observed as animals progressed through a block.

## 5 Conclusion

Altogether, this study provides confirmation of past results which suggest mPFC involvement in working memory processes. This study adds to our knowledge by causally revealing mPFC involvement in each working memory process (encoding, maintenance, retrieval) specifically when faced with decisions involving conflicting recent memories which require the flexible use of memories. Importantly, our data revealed a decrease in the number of reversals completed in session during mPFC return epoch disruption, an increase in regressive errors during mPFC choice epoch disruption and, an increase in perseverative errors resulting from delay epoch disruption. These epoch-selective increases in errors are suggestive of distinct working memory processes (encoding, maintenance, and retrieval) being interrupted as a result of mPFC disruption. Our VTE analysis revealed the mPFC is responsible for modulating this behavior as new information needs to be flexibly incorporated into working memory. Overall, our results fit a model in which the mPFC is required for retrieving context appropriate memories during decision-making but only becomes necessary to encode and maintain task relevant information in working memory during flexible decision-making. In this way, the mPFC becomes more engaged in processing working memory information as task demands change and uncertainty increases.

