## Supplemental Figure for "The medial prefrontal cortex during flexible decisions: Evidence for its role in distinct working memory processes"

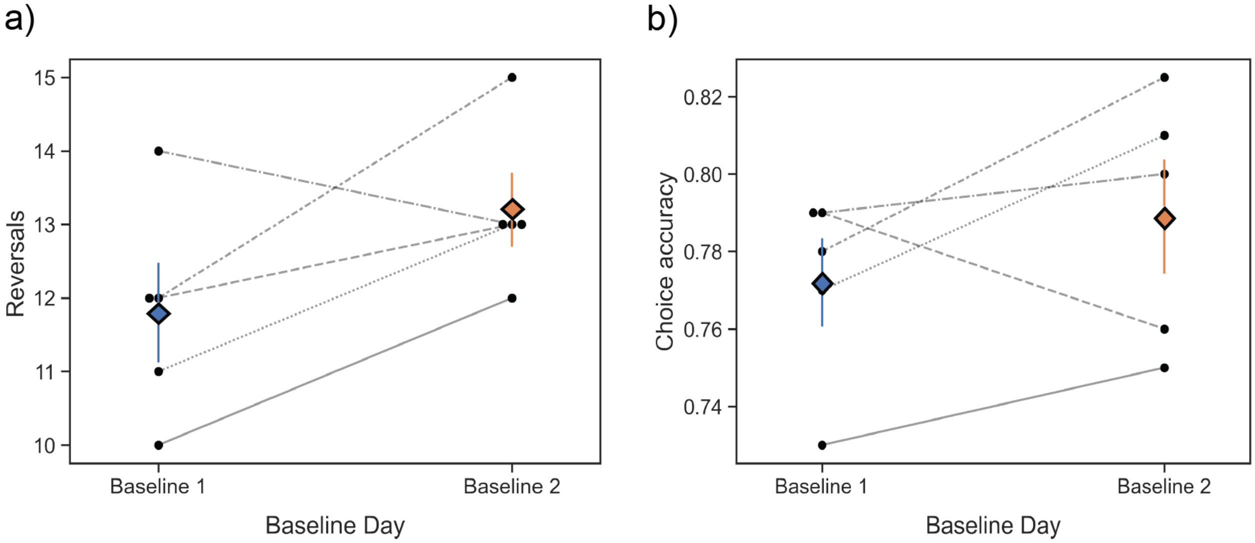


**Supplemental figure S1**

We compared performance metrics across Baseline Days 1 and 2 to check for the possibility that our optogenetic disruption in the mPFC caused performance deficits due to repeated mPFC stimulation. a) The number of reversals completed by animals on Baseline Day 1 vs Baseline Day 2 were not significantly different ($p$>.05). b) Choice accuracy did not significantly change ($p$>.05) from Baseline Day 1 to Baseline Day 2.
